# Three-Dimensional Reconstructions of Mouse Circumvallate Taste Buds Using Serial Blockface Scanning Electron Microscopy: I. Cell Types and the Apical Region of the Taste Bud

**DOI:** 10.1101/610410

**Authors:** Ruibiao Yang, Yannick K. Dzowo, Courtney E. Wilson, Rae L. Russell, Grahame J. Kidd, Ernesto Salcedo, Robert S. Lasher, John C. Kinnamon, Thomas E. Finger

## Abstract

Taste buds comprise four types of taste cells: 3 mature, elongate types: Type I, Type II, Type III; and basally-situated, immature post-mitotic Type IV cells. We employed serial blockface scanning electron microscopy to delineate the characteristics and interrelationships of the taste cells in the circumvallate papillae of adult mice. Type I cells have an indented, elongate nucleus with invaginations, folded plasma membrane, and multiple apical microvilli in the taste pore. Type I microvilli may be either restricted to the bottom of the pore or extend outward reaching midway up into the taste pore. Type II cells (aka receptor cells) are characterized by a large round or oval nucleus, a single apical microvillus extending through the taste pore, and specialized “atypical” mitochondria at functional points of contact with nerve fibers. Type III cells (aka “synaptic cells”) are elongate with an indented nucleus, possess a single, apical microvillus extending through the taste pore and are characterized by a small accumulation of synaptic vesicles at points of contact with nerve fibers. About one-quarter of Type III cells also exhibit an atypical mitochondrion amidst the presynaptic vesicle clusters at the synapse. Type IV cells (non-proliferative “basal cells”) have a nucleus in the lower quarter of the taste bud but have a foot process extending to the basement membrane often contacting nerve processes along the way. Type I cells represent just over 50% of the population, whereas Type II, Type III, and Type IV (basal cells) represent 19%, 15%, and 14% respectively.

## Introduction

In mammals, taste buds, the sensory end organs for the gustatory sense, comprise 50-100 spindle-shaped cells arranged like cloves in a garlic bulb. The apices of the taste cells extend into the taste pore, an opening in the epithelium, allowing interaction of tastants with the taste receptors situated on these apical extensions. The detailed ultrastructure of these and other cellular features of taste buds began with studies on rabbits (Engstrom and Rytzner 1956, Engstrom and Rytzner 1956) with the first ultrastructural studies on rodents being performed on the rat a decade later (Farbman 1965) and Gray & Watkins (Gray and Watkins 1965) and subsequently on the mouse, starting in 1974 (Mattern and Paran 1974). These studies converge on a common understanding of cell types and general features of mammalian taste buds (Kinnamon and Yang 2008).

Farbman (Farbman 1965) introduced the commonly adopted nomenclature for taste cells, including: peripheral (or edge), dark (Type I), light (Type II) and basal (Type IV) types. These so-called basal (Type IV) cells of taste buds are post-mitotic and despite the similarity in name, should not be confused with the proliferative basal cells that underlie all epithelia. Ultrastructural defining features of the various taste bud cells have included nuclear size and shape, overall configuration of cellular processes (Pumplin, Yu et al. 1997), apical microstructure (Kinnamon and Yang 2008) and relationship to nerve fibers and other cells in the taste bud (Kinnamon, Taylor et al. 1985). A distinct Type III cell, marked by readily-identifiable synapses onto nerve fibers, was added to the original dark-light scheme (Takeda and Hoshino 1975).

More recent studies show a correlation between functional-molecular features of taste cells and the older ultrastructural classifications. Type I cells exhibit many features of astrocytes including fine processes extending between the other cell types (Pumplin, Yu et al. 1997), neurotransmitter reuptake (Lawton, Furness et al. 2000) or breakdown (Bartel, Sullivan et al. 2006), and K^+^homeostasis (Dvoryanchikov, Sinclair et al. 2009). Type II cells express the receptors for bitter, sweet or umami (Chaudhari and Roper 2010) along with the downstream transduction components including PLCβ2, IP_3_R_3_ and TrpM5. Type II cells form unique synapses with nerve fibers involving large mitochondria with tubular cristae (so-called “atypical” mitochondria) closely apposed to the CALHM1/3 ATP release channels in the cell membrane (Taruno, Vingtdeux et al. 2013, Ma, Taruno et al. 2018, Romanov, Lasher et al. 2018). Strong depolarization of the Type II cells gates open the CALHM1/3 channels to release the neurotransmitter, ATP, into extracellular space. In contrast, Type III cells, which mediate sour, and possibly some portion of salty taste form typical chemical synapses with nerve fibers including a small accumulation of presynaptic vesicles in the presynaptic compartment (Kinnamon, Taylor et al. 1985). Both Type II and Type III cells are more regular, spindle shaped cells than are Type I cells which have diaphanous processes wrapping around other cells and nerve processes within the bud (Pumplin, Yu et al. 1997). Type IV cells express sonic hedgehog (Shh) and are recently generated, postmitotic cells with nuclei lying in the bottom half of the taste bud and capable of differentiating into any of the 3 elongate taste cell types (Miura, Kato et al. 2004, Miura, Scott et al. 2014). Other details regarding the morphology and interrelationships of these basal cells is unknown.

In the present study we have used serial blockface scanning electron microscopy (sbfSEM) to generate three-dimensional (3-D) reconstructions of longitudinal serial sections taken through circumvallate taste buds of mice. The sbfSEM technique, which entails large field, high resolution imaging of tissues prepared for electron microscopy became practical for routine use only in the last decade (Denk and Horstmann 2004). Previously, laborious reconstructions from serial sections of rodent and rabbit taste buds imaged with transmission electron microscopy were carried out by J Kinnamon and co-workers (Kinnamon, Sherman et al. 1988). In contrast, sbfSEM, where the imaging and serial sectioning are done automatically, permits acquisition of very large image stacks with better than 10 nm resolution encompassing nearly complete taste buds. This permits a more quantitative and complete analysis of ultrastructural features and classifications of cell types than with the previous technology. This paper focuses on the overall structure of taste buds and the constituent cell populations. Later papers will concern the interactions between cells and their relationships with nerve fibers.

## Materials and Methods

### Serial Blockface Scanning Electron Microscopy (sbfSEM)

Mice (C57/Blk6) were anesthetized with Fatal-Plus Solution® and perfused through the heart with 0.1% NaNO_2_, 0.9% NaCl, and 200 units sodium heparin in 100 ml 0.1 M phosphate buffer (pH7.3), at 35 °C to clear the circulatory system. The mice were then perfusion fixed with 2.5% glutaraldehyde and 2% formaldehyde containing 2mM CaCl_2_ in 0.025 M sodium cacodylate buffer (pH 7.3) at 35 °C for 10 minutes. Tongues were then removed and placed in the same fixative for 2-3 hours at 4 °C. The circumvallate papillae then were sliced into 200 µm thick vibratome sections.

For conventional transmission electron microscopy, a subset of sections were rinsed in buffer, then reacted with 2% Osmium Tetroxide in 0.05 M Sodium Cacodylate Buffer for 30 min. After three ddH_2_O rinses, the sections were placed overnight in 1% uranyl acetate in ddH_2_O and then stained *en bloc* in Walton’s lead aspartate at 60 °C for 40 min prior to embedding in Luft’s Epon.

Sections (200 µm thick) for serial blockface scanning electron microscopy (sbfSEM) were washed with 0.025 M cacodylate buffer (pH7.3) with 2 mM CaCl2, then incubated for 1 hour at 0° C in a solution containing 3% K4[Fe(CN)6] in 0.025M cacodylate buffer pH7.3 with 2mM CaCl_2_ combined with an equal volume of 4% aqueous OsO_4_. After the first heavy metal incubation, the sections were washed with H_2_O at room temperature 5×3 min. and then placed in 1% thiocarbodhydrazide solution for 20 min at room temperature. After washing, the sections were placed in 2% OsO_4_ for 30 min at room temperature. Following this second exposure to osmium, the tissues were washed in H_2_O 5×3 min at room temperature, then placed in 1% UO_2_(OCOCH_3_)·2H_2_O at 4 °C overnight. The next day, the tissues were stained *en bloc* with Walton’s lead aspartate for 30 min at 60 °C in 0.066 g of Pb(NO_3_)_2_ in 10 ml of aspartic acid stock and pH adjusted to 5.5 with 1N KOH. Sections were then dehydrated using an increasing series of ice-cold alcohol solutions before transferring to propylene oxide 3×5 min. and final embedment in Lufts Epon 3:7 at 60 °C overnight.

Semithin sections of the tissue blocks were examined to identify regions containing taste buds. The blocks then were trimmed and mounted on an aluminum pin, coated with colloidal silver paste around the block edges, and then examined with a Zeiss Sigma VP system equipped with a Gatan 3View in-chamber ultramicrotome stage with low-kV backscattered electron detectors optimized for 3View systems. Areas of the blockface containing taste buds were identified and then these regions were imaged routinely at 2.25 kV, at 5-10 nm/pixel resolution (30 µm aperture, high current mode, high vacuum), with field sizes between 80-250 µm in x,y and approximately 500 slices with 70-85nm thickness were generated. The resulting image stacks are aligned in Image J and montaged in Photoshop (Adobe Systems). Segmentation and reconstruction was carried out using *Reconstruct* software (Fiala 2005). Each composite image was viewed separately and cell membranes, nuclei, etc., were segmented using the pencil feature. Segmentations from each image for each structure were combined to create 3D rendered images in *Reconstruct*.

## Results

### General Features

A longitudinal section through a circumvallate taste bud shows a prominent taste pore (TP) with examples of diverse apical processes from Type I, Type II and Type III cells (Figure 1). This taste bud contains approximately 50 taste cells. The boundaries of the taste bud – both lateral and basally are somewhat arbitrary in terms of whether to include edge cells and cells situated basally along the basal lamina. We have taken a fairly conservative approach, including in our analysis only cells that are clearly within the confines of the taste bud and interacting with other cells and nerve fibers in the bud. In Fig. 1, the cells lying along the basement membrane are not included in our analysis in that the cells lie entirely below the taste bud proper and have no apically directed processes extending within the taste bud.

**Figure 1.**
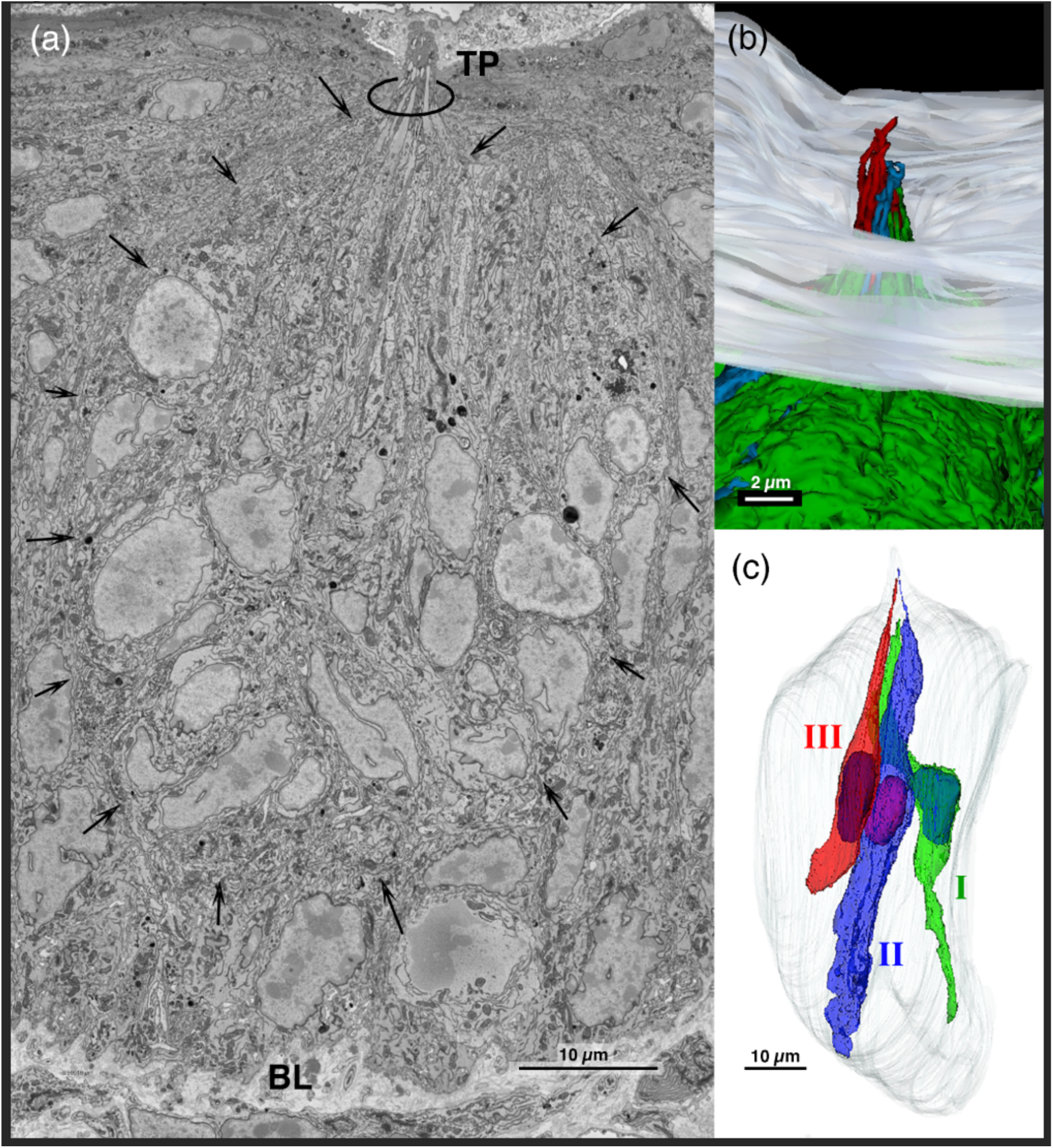
**(a).** A single plane, low magnification micrograph from the sbfSEM dataset showing a mouse circumvallate taste bud from basal lamina (BL) to a taste pore (TP). A taste bud contains about 50 taste cells, some with a large, circular nucleus (Type II cells), others with irregular, elongate nuclei with nuclear invaginations (Type I or III cells). **(b).** 3-D reconstruction of a taste bud showing the three different cell types in relation to the surface of the epithelium (gray). Type I cell in green; Type II cell in blue, and Type III cell in red. Note that the Type II and III cells each extend a single microvillus farther into the taste pore than do the Type I cells. **(c).** 3-D reconstruction showing 3 taste cells, one of each type within a single taste bud.

Even at this low magnification, diverse cell types are evident according to nuclear and cytoplasmic features, e.g. cytoplasmic organelles vary and some nuclei are fairly smooth and round while others are crenulated. With conventional transmission electron microscopy, Type I cells have an electron-dense cytoplasm while Type II cells have an electron-lucent cytoplasm. The tissue preparation procedures for sbfSEM, however, leave the electron density of the different cell types less distinctive. This makes classification of the different cell types based on electron density problematic. Hence, we distinguish the different cell types primarily based on cell shape, organelles, nuclear morphology and apical processes. We further note the presence of conventional synapses or atypical mitochondria at points of contact between the taste cells and nerve fibers. All three cell types have apical processes that may extend into the taste pore, but Type I apical processes do not extend as far into the taste pore (Figures 1b, c; 2a, b) (also see below) as the other 2 elongate cell types.

**Figure 2.**
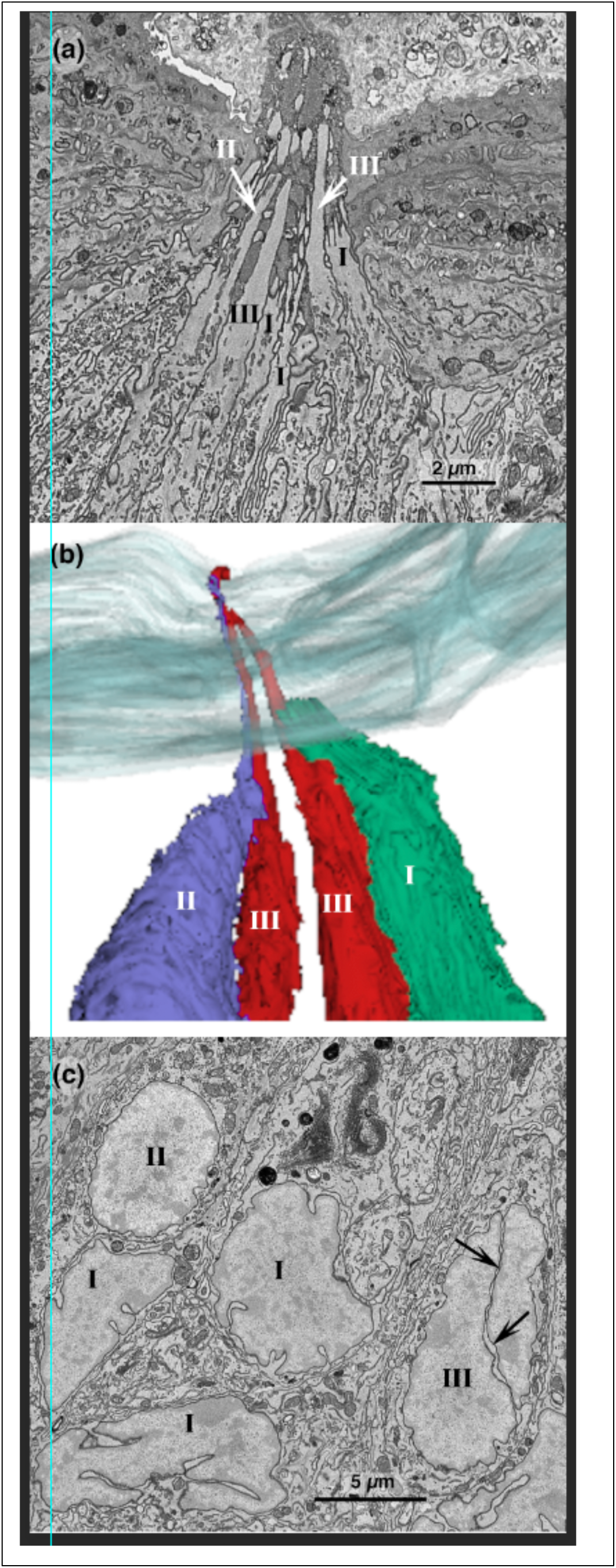
The different cell types are distinguishable by relationship to the taste pore and nuclear structure**. (a):** High magnification of the taste pore region showing some taste cells with single thick microvilli (Types II, III) extending high into the pore while others (Type I cells) extend branched microvilli ending in a bushy form near the base of the pore region. **(b):** Four taste cells from this taste pore rendered as individual objects showing the difference in height of apical extensions of the different cell types. **(c)** Single plane image through a taste bud showing the different morphologies of cell nuclei of the different cell types. Type I cell nuclei are irregular and often exhibit numerous invaginations of the nuclear envelope. Type II cell nuclei are relatively round or oblate with a smooth nuclear envelope. Type III cell nuclei are elongate and show one or more deep invaginations (arrow) of the nuclear membrane.

## Distinguishing Features of Cell Types

### Type I Cells

The characteristic morphology of Type I cells is shown in Figures 2 & 3 and includes an irregular, indented nucleus, refolded lamellate processes enwrapping nerve fibers and other cells, and prominent Golgi apparatus. The Type I cell nuclei are generally elongate and may possess prominent invaginations, which can make distinguishing Type I cells from Type III cells challenging (Figure 2c). A hallmark of Type I cells is the presence of thin, lamellate sheets of membrane that envelop other taste cells or nerve processes (Figure 4a). Although Type I taste cells come into extensive, intimate contact with nerve processes, we have not observed any distinctive organelles – either vesicles or mitochondria -- at points of contact between Type I cells and nerve processes or other taste cells. In many cases, the nerve process indents the cell in the region of the nucleus leaving scant cytoplasm in the space between the nerve fiber and the nucleus (Fig. 4). In some of these cases, either particles or strands of electron dense material may lie in the narrow cytoplasmic space between the cell plasma membrane covering the nerve process and the nuclear membrane.

**Figure 3.**
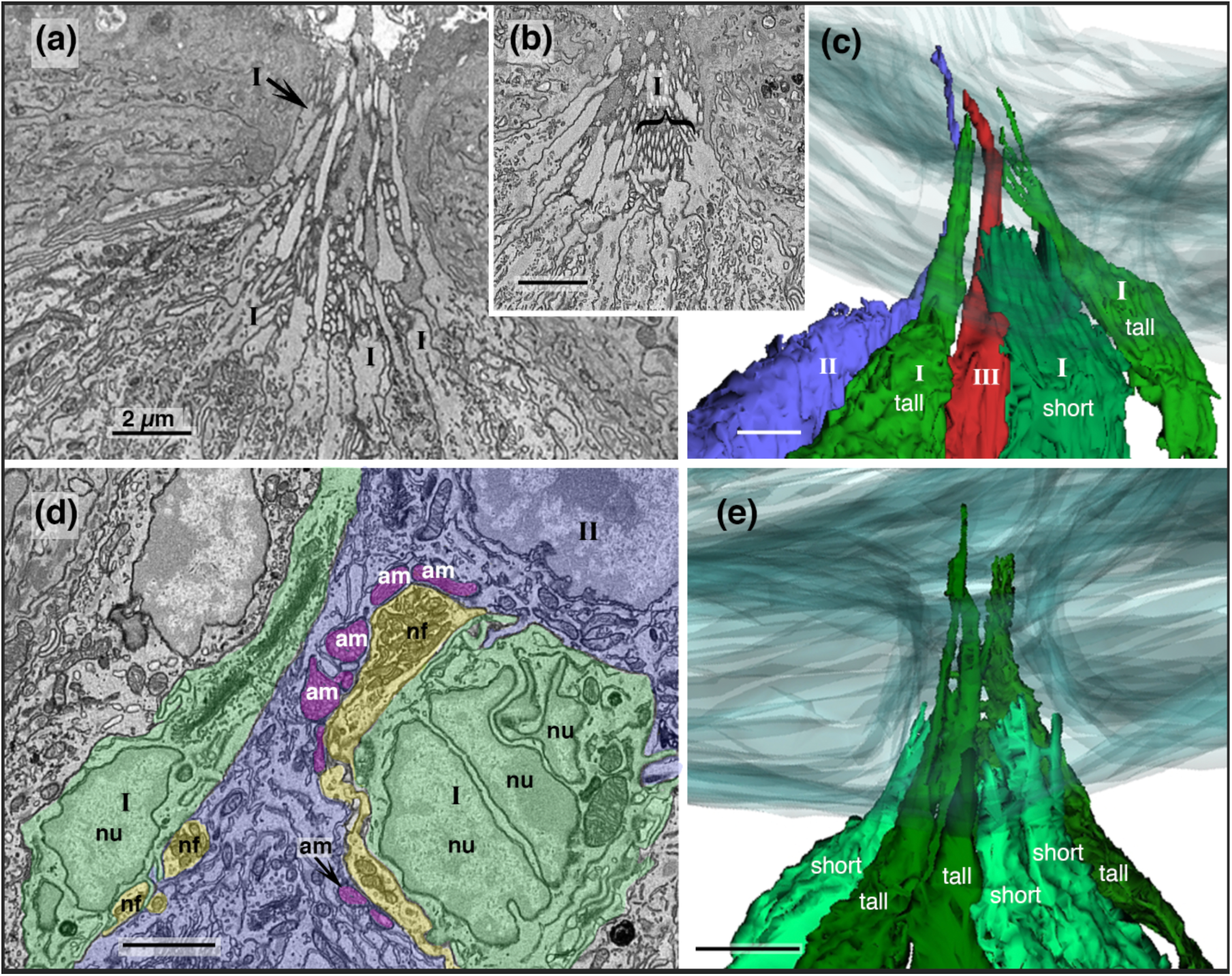
Type I cell morphological features. **(a, b, c, d)** Some Type I cells end in an apical tuft in or just below the base of the taste pore while other type I cells extend a branched process outward into the taste pore but rarely reaching outward as far as Type II or Type III cell processes. **(a)** Single plane image of the taste pore region. Two types of apical specializations are evident for Type I cells: bushy and elongate. The apical processes of tall Type I cells are stout and extend upwards into the taste pore emitting several thick branches along the way. In contrast, short-mv Type I cells end below, or extend only a short distance into, the taste pore where they end as a terminal bush of short, thin microvilli. **(b)** Apical pore region showing the bushy type, fine microvillous processes of several Type I cells (bracket). **(c)** 3D reconstruction of three Type I cells (green) along with a Type III cell (red) and a Type II cell (blue) for comparison. The tall Type I cells are shorter than any of the Type II and Type III cells although they are taller than the bushy, short-mv Type I cells. **(d)** The Type I cells (green) exhibit an irregular nucleus (nu) with nuclear invaginations. A common characteristic of Type I cells is that the membrane extends lamellate processes to envelop nerve fibers (yellow) or other taste cells (Type II cell, gray blue with atypical mitochondria (purple) at point of contact with the nerve fiber). Despite close contact with nerve fibers (nf, yellow), Type I cells lack evidence for synaptic specializations, e.g. vesicles or mitochondria, at points of contact. **(e)** 3-D reconstruction of six Type I cells in a single taste bud showing the two different apical morphologies. Short cells (light green) end in a busy apical microvillus whereas tall Type I cells (dark green) have a longer branched apical process.

**Fig 4:**
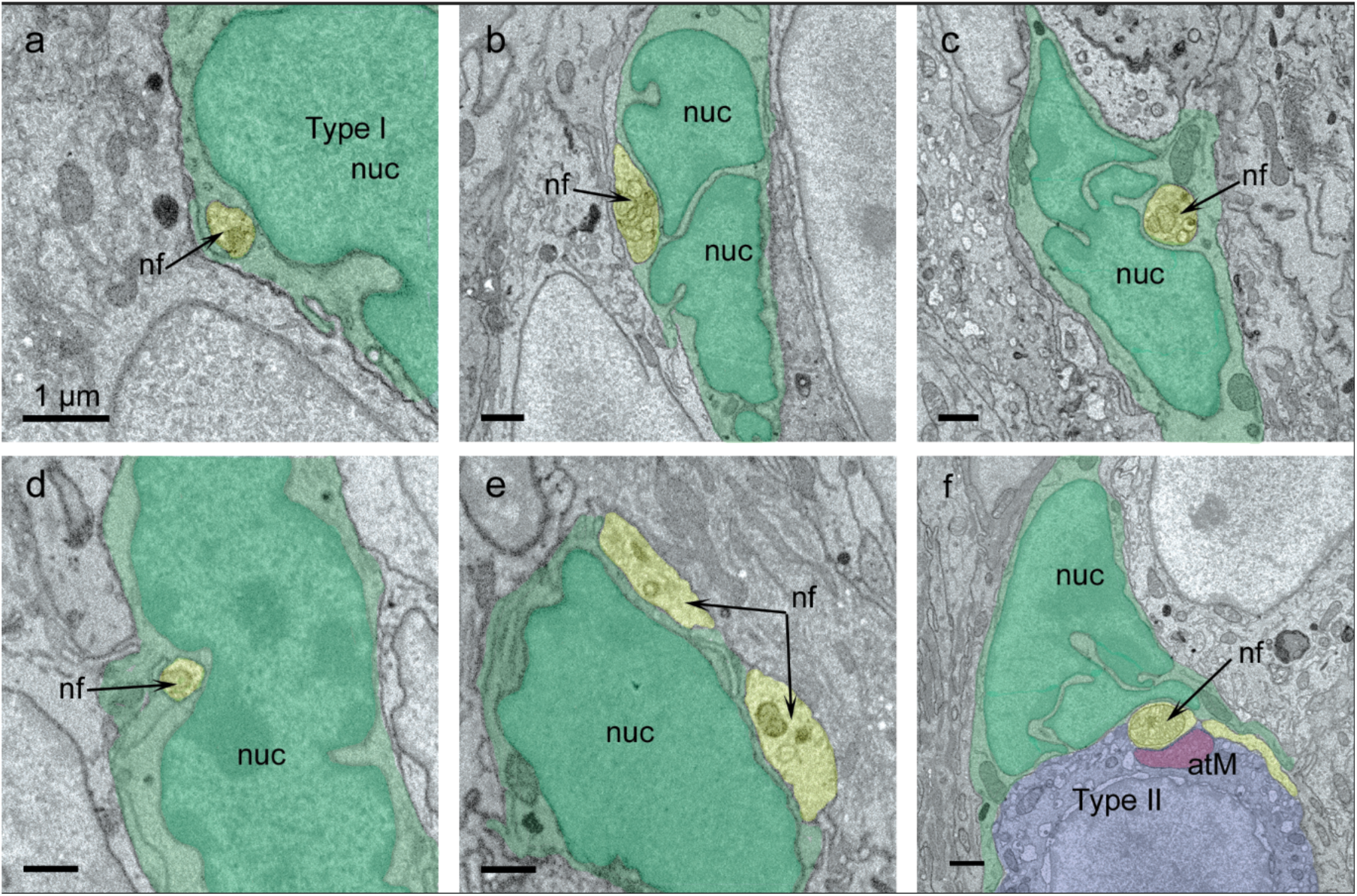
The nucleus (nuc) of Type I cells (green) often lies in close association with nerve fibers (nf; yellow). In many cases, the Type I cells fully embrace **(a, c, d)** nerve fibers. Despite the intimate association of the nerve fibers and Type I cell nuclei, no clear membrane thickenings or synaptic specializations are evident at the point of contact.

The apical processes of Type I cells appear to be of two different morphologies, possessing either: a single, tall microvillus (tall-mv subtype) or numerous short, bushy microvilli (short-mv subtype). The tall-mv Type I cell has a stout microvillus which extends at least midway up into the taste pore. This apical microvillus often exhibits short lateral extensions, either at its root near the bottom of the taste pore or as short branches along the trunk of the process, giving it a somewhat shaggy appearance. These elongate, Type I cells also exhibit two slightly different morphologies. In one subtype the long microvillous arises simply from the apex of the cell while in the other subtype, the long microvillus arises from a small collection of short protuberances near the base of the taste pore. The short-mv Type I cell has numerous apical microvilli which appear as a bushy terminus, reaching upward a short distance forming the floor of the taste pore, rarely reaching even midway up into the taste pore. The microvilli of the short-mv Type I cells do not have thick branches like those of the tall-mv Type I cells. Instead, the short-mv Type I cell microvilli terminate in an expanded bush-like cluster. A 3-D rendering of the two Type I cell apical specializations is shown in Figure 3c, e.

Some Type I cells do not extend outward as far as the floor of the taste pore although they lie entirely within the imaged volume. Despite the absence of an apical microvillus, these Type I cells exhibit the other characteristics of mature Type I cells including indented nucleus, prominent Golgi apparatus, and lamellate processes wrapping nerve processes and other cell types.

### Type II Cells

The characteristic features of Type II cells are round, smooth nuclei and a single, thick apical microvillus that extends well into the taste pore, often reaching the level of the surface of the surrounding epithelium (Figure 1b, 5a). This contrasts with Type I microvilli, which extend only part way into the taste pore. A common feature of Type II cells is a large, round nucleus with a predominately homogeneous nuclear matrix, although patches of heterochromatin may be present, especially adherent to the nuclear membrane (Figure 2c; 5b, c). Some, but not all Type II cells contain atypical mitochondria that occur exclusively at the close appositions between the Type II cell and a nerve process (Romanov, Lasher et al. 2018). Whereas typical mitochondria usually contain lamellar cristae arranged in a shelf-like configuration, atypical mitochondria contain tubular cristae with no apparent organization (Figure 5d).

**Figure 5.**
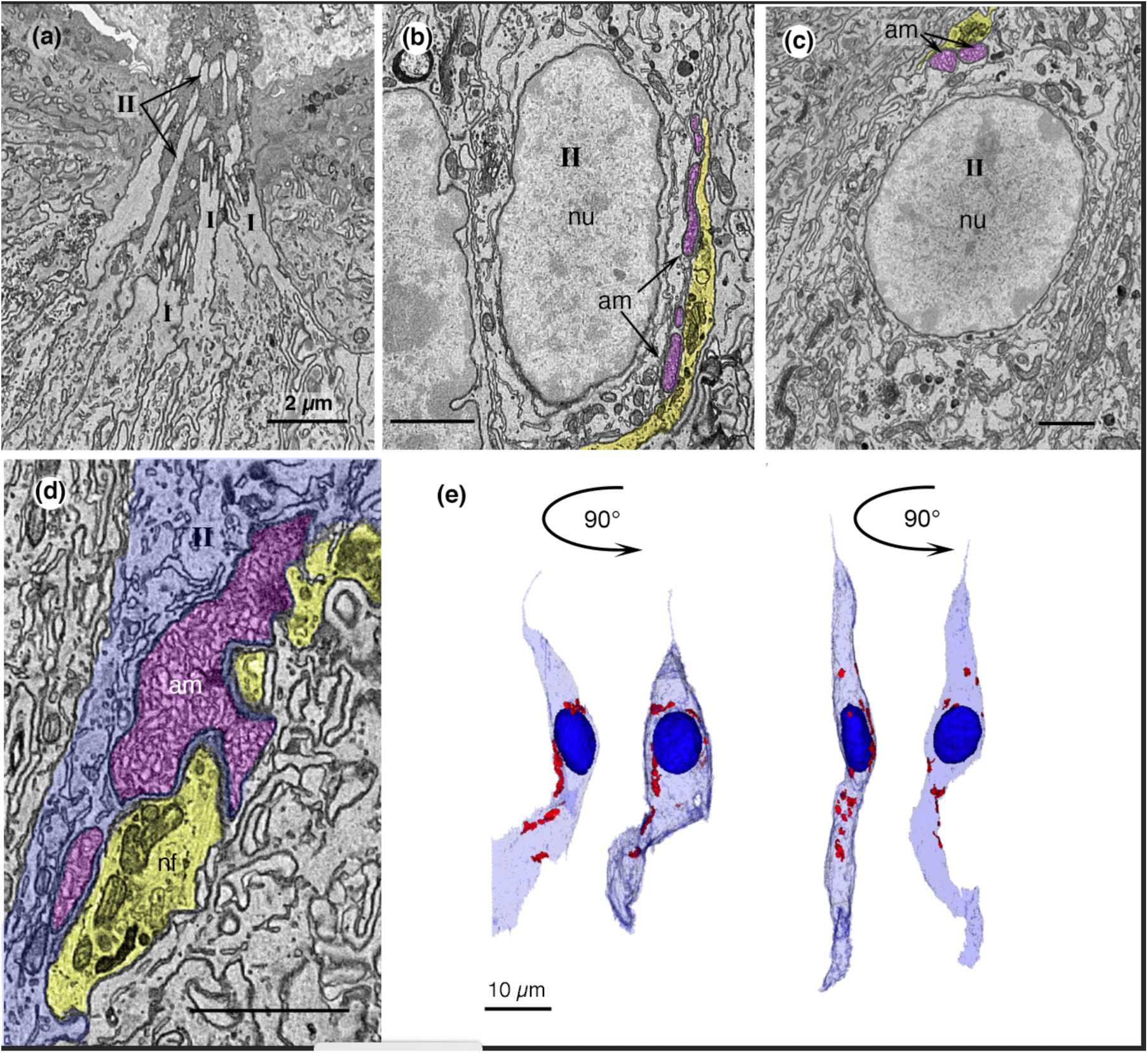
Morphological features of Type II cells. **(a)** Type II taste cells extend a single thick apical microvillus (II) high into the taste pore. **(b, c)** Type II cells have a large round nucleus (Nu) with relatively sparse heterochromatin, and electron lucent cytoplasm. The nucleus may appear either circular or ovoid. At some points of contact with nerve fibers (yellow), atypical mitochondria (am; red) are evidence of synaptic specialization (see Romanov et al 2018). **(d)** Higher magnification showing a large atypical mitochondrion (red) abutting the membrane facing the nerve fiber (yellow). **(e)** Reconstructions of 2 Type II cells with each cell showing its different appearance when rotated 90°. When viewed from different perspectives, the nucleus (dark blue) of a Type II cell can appear elongate (left side of each pair) or circular (right half of each pair). Red spots indicate the position of atypical mitochondria in each cell. Taste pore up.

### Type III Cells

Type III cells have a single, long, blunt microvillus that extends, like Type II cell microvilli, well into the taste pore (Figures 1b; 2b and 6a) often reaching to or even above the surface level of the surrounding epithelium. In general, the microvillus of a Type III cell extends farther outward than the microvilli of Type II cells in the same taste bud. Type III cells display an elongate nucleus with prominent, deep invaginations in the nuclear membrane (Fig. 2c, 5b). The most distinguishing feature of Type III cells is the presence of conventional synapses with a small accumulation of presynaptic vesicles. These synapses are characterized by the presence of 40-60 nm clear vesicles, some of which appose the presynaptic membrane (Figure 6b, s1, s2) abutting a nerve process. In conventional transmission electron micrographs, the presynaptic membranes are thickened and electron-dense; this is not as apparent with the sbfSEM technique.

**Fig. 6.**
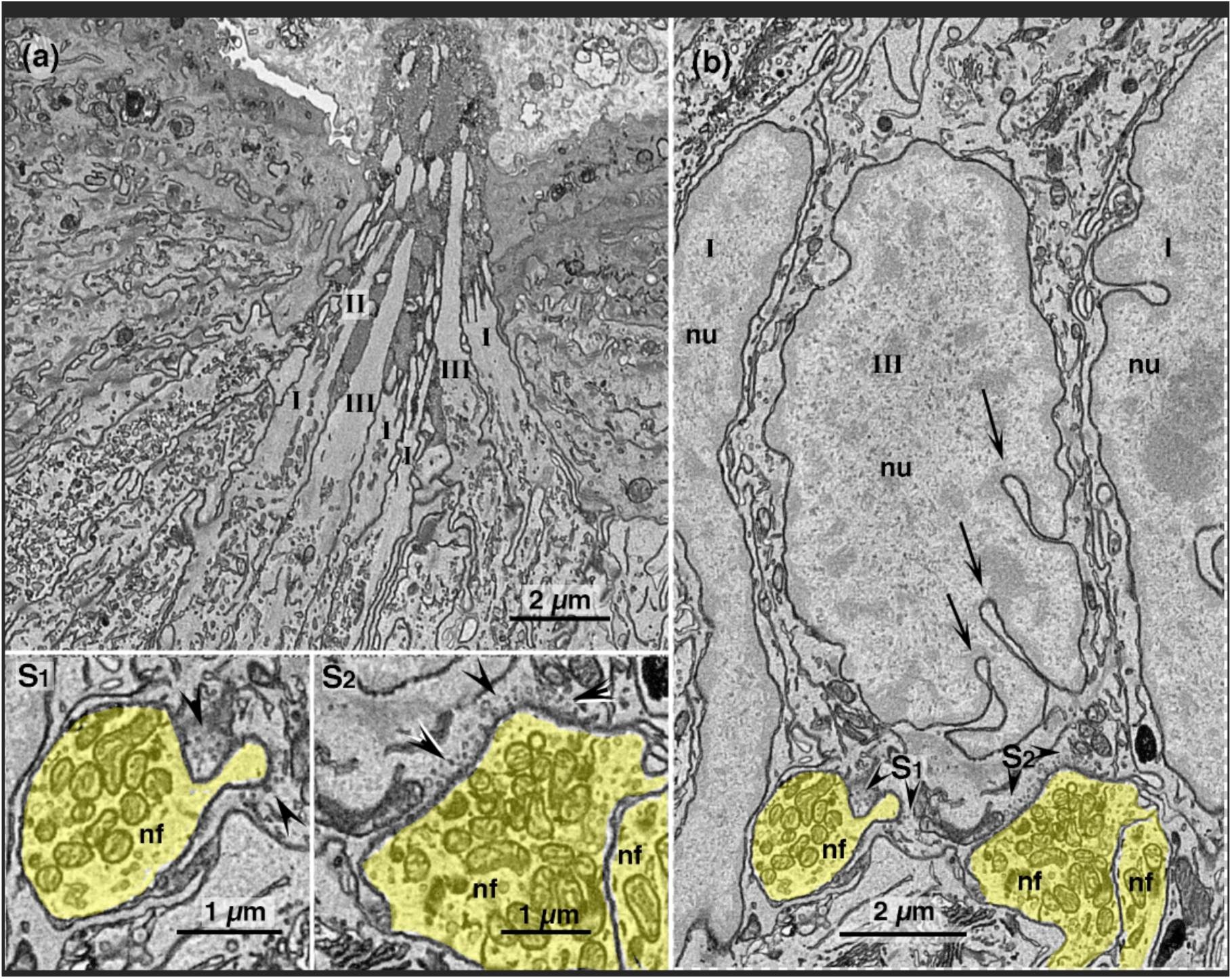
Morphological features of Type III cells. **(a)** Type III cells are slender extending a single, large, and blunt microvillus (III) into the taste pore. Note that the Type III cell microvillus is thicker than Type II cell (II) microvilli. **b.** Type III cell nuclei are irregular and elongated, often with deep invaginations (arrows). Also visible are two adjacent Type I cell nuclei (nu) with small invaginations. Classical chemical synapses (S1, S2 enlarged at lower left) with synaptic vesicles (arrow heads) appear at some points of contact between the Type III cell and nerve fibers (nf, yellow). **S1, S2:** Enlargements from panel (b) showing several clear, synaptic vesicles (arrow heads) at points of contact with the nerve fibers.

In our examination of 23 Type III taste cells, we have identified 5 (21.7%) that also exhibited one or more large mitochondria with tubular cristae at a point of contact with a nerve fiber that also shows an accumulation of presynaptic vesicles (Fig. 7) although not all synaptic contacts of these cells exhibit an atypical mitochondrion. These cells have the characteristic features of a Type III cell, i.e. indented nucleus and single long apical microvillus as well as an accumulation of vesicles where the cell membrane contacts a neural process. The mitochondria at the points of vesicle accumulation appear identical to the “atypical mitochondria” associated with neural contacts of Type II cells. Both types of mitochondria are larger than the other mitochondria in the cell and have irregular, tubular cristae not common in other mitochondria of the same cell.

**Fig. 7:**
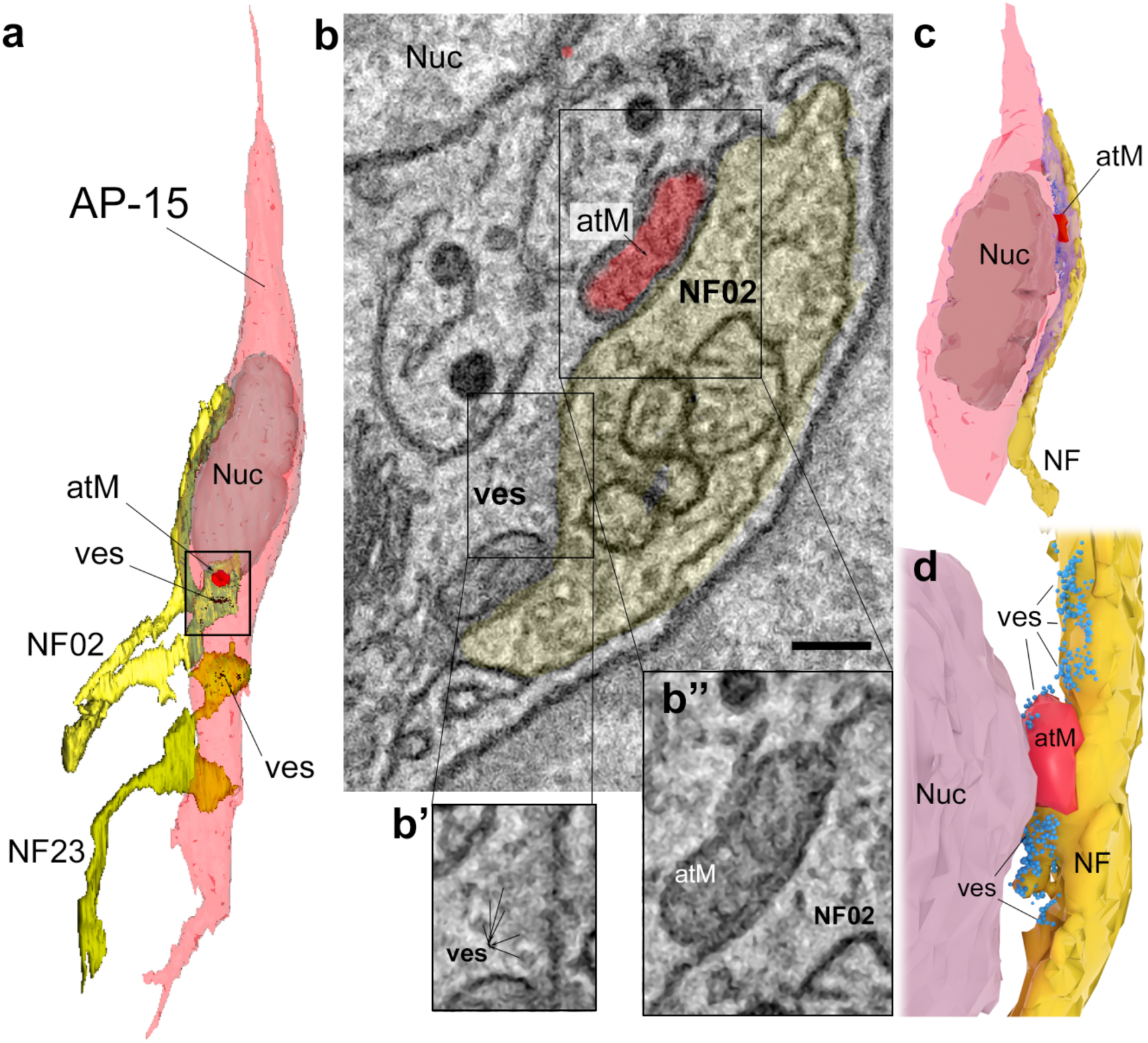
Some Type III cells exhibit mitochondria with irregular cristae associated with vesicle clusters at points of contact between the Type III cell and nerve fibers. **a**. Reconstruction of a Type III taste cell (AP-15) and 2 associated nerve fibers (NF02 and NF23). Accumulations of vesicles (ves) near the point of cell-nerve contact are indicative of Type III cell synapses. At one of these contacts, we also find one or more mitochondria (red) with irregular cristae closely apposed to the cell membrane. **b**. Single plane sbfSEM image of the region containing the mitochondrion of interest showing the location of vesicles (ves) and mitochondrion (atM) in relation to the nerve process (NF02). **b’.** Enlargement of the vesiclular accumulation framed by the box in **b**. This image was enlarged by 125% and filtered with a Photoshop Dust & Scratches Filter (3 pixel radius) to reduce graininess. **b”.** Enlargement of the boxed area in **b** around the mitochondrion processed similarly to **b’** to show irregular cristae. **c.** Reconstruction of another Type III cell (pink) with an irregular mitochondrion (red) at the point of contact with a nerve process (yellow). Higher magnification 3D rendering showing the relative position of vesicles (blue), mitochondrion (red) and underlying nerve process (yellow). For purposes of visualization the cell membrane of the Type III cell was rendered transparent.

### 3-D Reconstructions of the Taste Pore Region

Figures 1b and 2b show 3-D reconstruction of the microvilli from a taste bud extending through the extragemmal epithelial surface. Figures 2a, 3a, and 5a show low magnification images of the microvilli of two taste buds. A higher magnification 3-D reconstruction of one of the taste buds demonstrates how the microvilli of the Type II cells (blue) and the Type III cells (red) extend farther into the oral cavity than the apical microvilli of the Type I cell. Figure 3c shows the same taste bud without the surrounding epithelium and from a slightly different angle, again demonstrating how the Type II and Type III microvilli extend further into the oral cavity when compared with the Type I microvilli. A movie (Suppl. Data) of the same taste bud graphically shows the apical processes of the different cell types and the extent to which they project into the oral cavity.

### Type IV cells (“Basal” cells)

Type IV cells are characterized by a nucleus situated in the lower quarter of the taste bud (Fig. 8a) and lacking extensive apical process. Further, these cells do not display any of the characteristic features by which we could assign them into other cell classes. Type IV cells have a spheroidal irregular nucleus with scattered patches of heterochromatin and a relatively thin rim of cytoplasm surrounding the nucleus. These cells are elongate in the longitudinal axis of the taste bud and have a basal extension contacting the basal lamina (Fig. 8b) as well as the plexus of nerve fibers underlying the taste bud. Although basal cells contact nerve fibers, no apparent membrane thickenings or specialized contacts occur, including classical synapses with vesicles or contacts with atypical mitochondria. Although basal cells often abut one another or even other cell types in the bud, no apparent membrane specializations occur at these points of contact. Some Type IV cells have a short apical extension which may partially envelop surrounding elongate taste cells (Fig. 8b).

**Figure 8.**
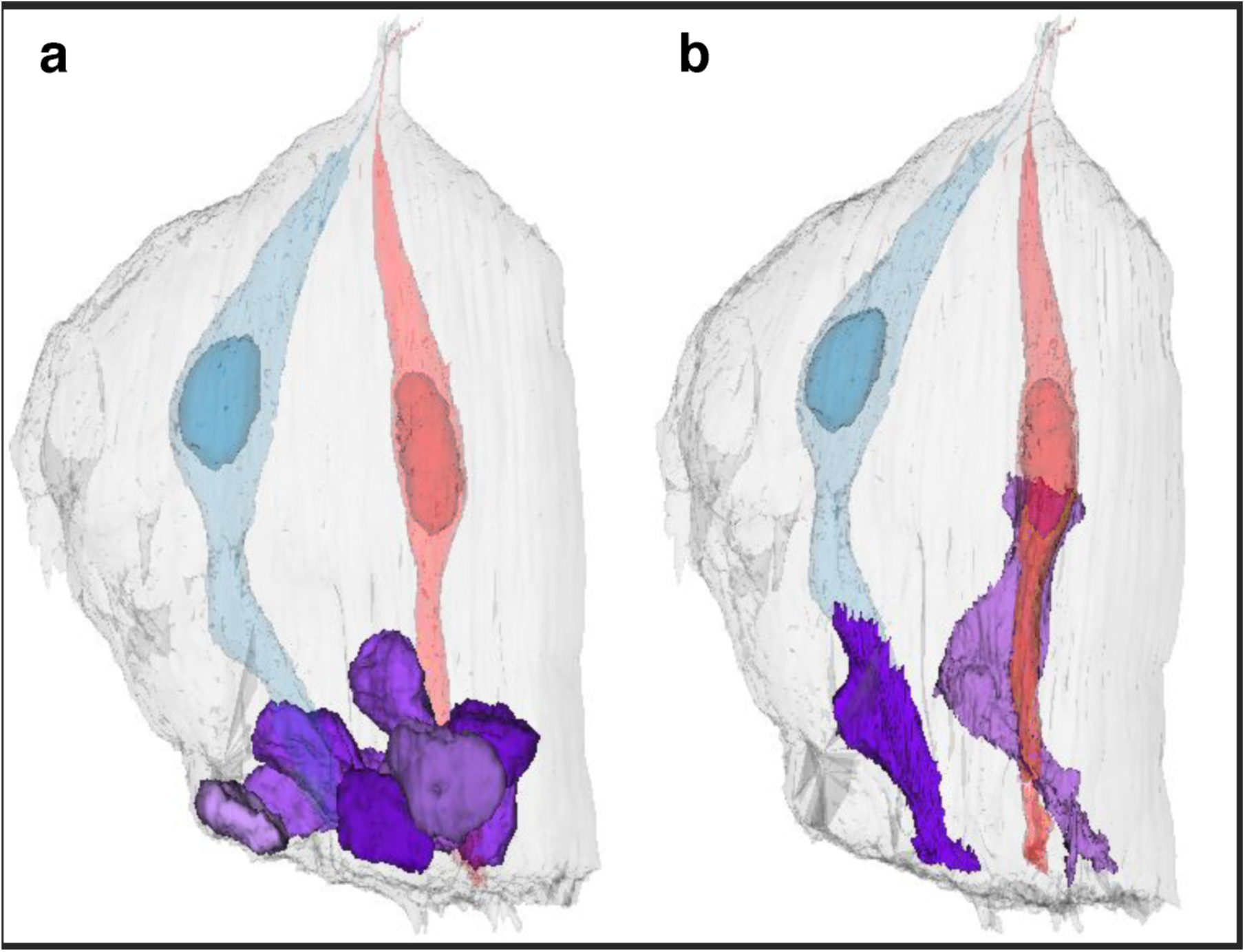
**a.** Type IV (basal) cells have nuclei (magenta) situated in the lower quarter of the taste bud while the nuclei of typical Type II (blue) and Type III (red) cells lie higher in the bud. **b.** The Type IV cells have a basal process extending to the basal lamina beneath the taste bud. Along the way, this process frequently contacts the basal plexus of nerve fibers penetrating the base of the bud. Some basal cells also have an apically directed process which may embrace one or more Type II or Type III cells.

### Cell Type Counts

Our data set includes 4 samples through incomplete taste buds from circumvallate papillae of 2 adult mice. These data allowed us to make population estimates of each cell type in each taste bud although we did not include any taste cells that extended outside of the boundaries of the data set. We fully segmented all Type II and Type III cells that extended into or nearly into the taste pore along with several Type I cells some of which did not reach the taste pore but nonetheless lay entirely within the segmented volume. In addition, we characterized by cell type without fully segmenting, all taste cells whose nuclei were fully contained within the sampling volume giving a total of 160 cells within 4 taste buds. The smallest sample included 13 cells from a small taste bud and 86 cells from the largest taste bud. Data for each of the 4 taste buds is given in Table 1. Taken together, the cell type counts show that Type I cells comprise 52.5% of the population; Type II cells 18.8%; Type III cells 15% and Type IV, 13.8%.

**Table 1:**
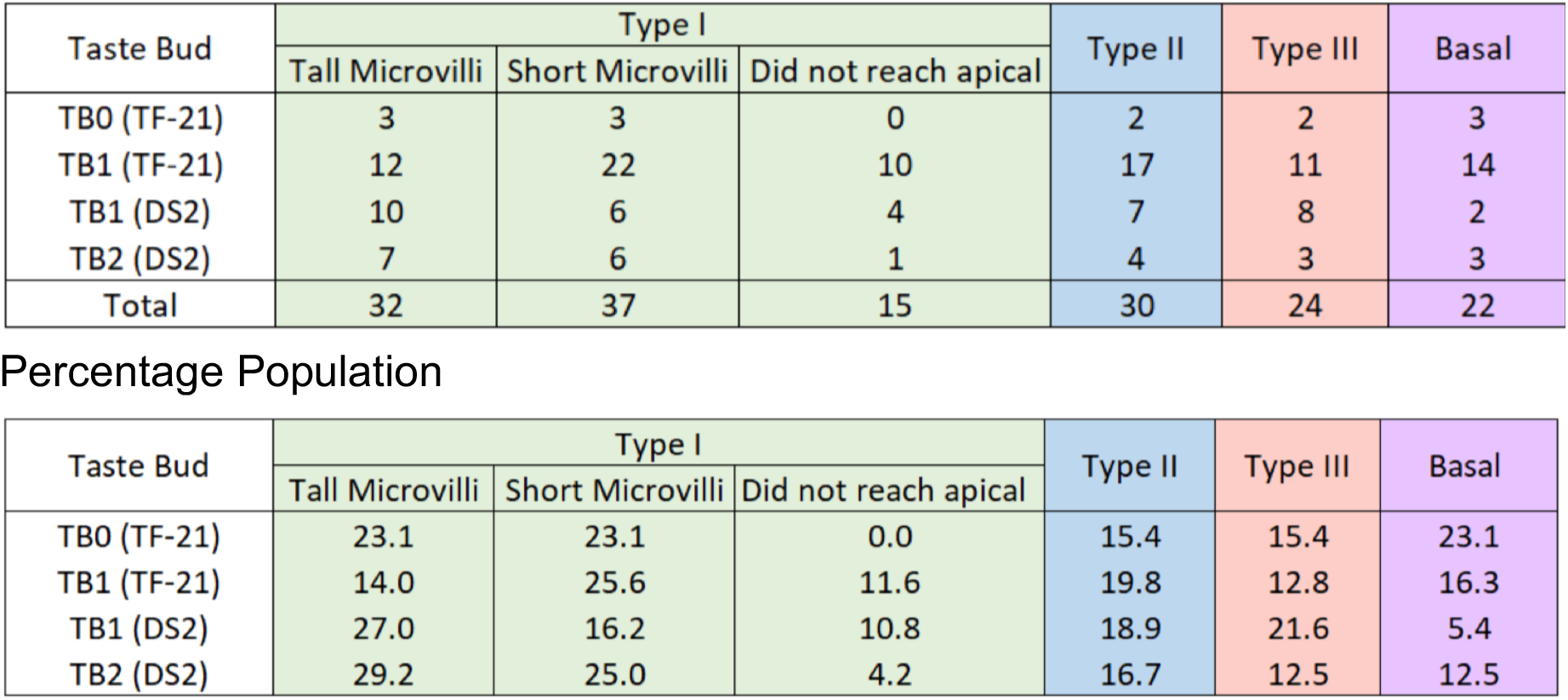
Counts (above) and population estimates (below) of different Types of Taste Cells in 4 taste buds of the Circumvallate Papilla.

## Discussion

The most commonly accepted contemporary delineation of mature taste cells divides them into 3 types of elongate, differentiated cells: Type I cells are considered glial-like, Type II cells underlie transduction of sweet, bitter and umami, forming unusual channel-based neurotransmitter release synapses with the afferent nerves ((Taruno, Vingtdeux et al. 2013, Ma, Taruno et al. 2018, Romanov, Lasher et al. 2018)) while Type III cells mediate sour taste and form classical synapses with synaptic vesicles at points of contact with nerve fibers. Our detailed analysis of the sbfSEM data confirm in broad terms this tripartite distinction between mature cell types.

These 3 morphological types of mature taste cells correlate well, but not exactly, with expression of immunochemical markers, whose expression has been used to estimate populations of different cell types in taste buds {Ma, 2007 #2283;Ohtubo, 2011 #1986}. In taste buds of fungiform papillae of mice {Ma, 2007 #2283}, Type I cells are the most common cell type, representing over 50% of the total population, with Type II cells next most abundant, (30-38%%) and Type III cells described as relatively rare, about 5-6% (Chaudhari and Roper 2010, Ohtubo and Yoshii 2011). The proportions are somewhat different in taste buds of murine circumvallate papillae which show substantially greater proportions of Type III cells (roughly 15%) with slightly smaller proportions of Type II cells (20%) (Ueda, Fujii et al. 2003, Ma, Yang et al. 2007). Our analysis is consistent with the result from circumvallate papillae reported by Ma et al {Ma, 2007 #2283}, i.e. that Type III cells are somewhat less abundant than Type II cells but still represent about 15% of the population. In contrast, taste buds of fungiform papillae have a much lower proportion of Type III cells.

The most striking results of this study are: 1) the presence of multiple subtypes of Type I cells distinguished by differences in apical microstructure and degree of extension toward the taste pore, 2) differences in structure of Type II taste cell apices between mouse and rats, 3) the presence of an atypical mitochondrion associated with presynaptic vesicle clusters in a significant proportion (21.7%) of Type III cells, and 4) the presence in Type IV cells, of a foot process that intertwines with gustatory nerve fibers and retains contact with the basal lamina.

### Type I cell subtypes

The 3D reconstructions reveal that some Type I cells end within the taste bud well below the taste pore region. The large majority of Type I cells extend to the taste pore but end apically in 2 different fashions: i) short, bushy microvilli ending near the floor of the taste pore (short-mv), and ii) longer, thick slightly branched microvilli that extend outwards more than half way up in the taste pore (tall-mv). The differences in apical microstructure of the different Type I cells may be related to the relatively rapid turnover of these cells in taste buds. The average lifespan of a Type I cell may be as short as 8 days (Perea-Martinez, Nagai et al. 2013), so at any time, 10-30% of Type I cells should be immature or senescent and apical microstructure may relate to the relative age of a Type I cell. For example, Type I cells not reaching the taste pore may be the youngest, while those with prominent long apical microvilli may be the most mature. Of course, other age-related schemes are possible. Furthermore, some Type I cells may live 24 days or longer (Perea-Martinez, Nagai et al. 2013) and it may be that shorter and longer-lived Type I cells display different apical microstructure.

Conversely, the existence of multiple morphological subtypes of Type I cells leads to speculation that there may be corresponding functional subtypes. The very short Type I cells that lack apical microvilli and do not approach the taste pore likely serve the glial-like functions usually attributed to Type I cells: degradation and reuptake of neurotransmitters, physical separation and isolation of other cell types, and ionic buffering (Pumplin, Yu et al. 1997, Bartel, Sullivan et al. 2006, Dvoryanchikov, Sinclair et al. 2009, Chaudhari and Roper 2010). In contrast, the Type I cells with apical microvilli reaching the taste pore might directly participate in taste transduction.

The taste pore is filled with a mucus substance that permits entry of small molecules such as lanthanum (Holland, Zampighi et al. 1991) but may prove a barrier to larger molecules such as sugars and many organic bitter compounds. The taller, branched microvilli within the taste pore likely have access to typical taste stimuli that could diffuse into the mucus substance filling the taste pore and so tall Type I cells could respond directly to such compounds. Small ionic taste substances such as salts and protons (sour), likely permeate well into the taste pore and therefore could directly interact with even the short-mv Type I cell apices Furthermore, these short-mv Type I cells, would be adequately acidified by any acids applied to the apical surface (Richter, Caicedo et al. 2003) and thus could participate in transduction of acids as well.

Although Type II and Type III cells are generally considered to be responsible for transduction of respectively, bitter, sweet and umami (Type II) and sour (Type III) (Chaudhari and Roper 2010), the possibility of Type I cells participating in transduction is not without precedent. (Vandenbeuch, Clapp et al. 2008) identified a population of taste cells expressing the amiloride-sensitive Na^+^channel potentially underlying some components of salt transduction, but that lacked the physiological hallmarks of both Type II and Type III cells, suggesting that Type I cells may respond to Na^+^ions. If a subset of Type I cells do participate in salt transduction, the question arises as to how the Type I cells might communicate with the nerve fibers. Although Type I cells have extensive areas of apposition to nerve fibers, the Type I cells exhibit no obvious synaptic specializations – either vesicular or mitochondrial – at these points of contact. Since all taste transmission requires activation of purinergic P2X receptors on the nerve fibers (Finger, Danilova et al. 2005, Vandenbeuch, Larson et al. 2015), the lack of presynaptic specializations in Type I cells suggests an unconventional means of transmitter release, e.g. by transporters or ion channels -- similar to the means by which astrocytes release gliotransmitters in the brain (Yoon and Lee 2014, Bang, Kim et al. 2016).

### Apical microstructure of taste cells

Our reconstructions of apical microvilli in mouse circumvallate taste buds reveal differences in microstructure compared to previous reports on cell apices in both rat and bovine taste buds (Yang, Tabata et al. 2000, Tabata, Wada et al. 2003). In circumvallate taste buds of both rats and cows, Type II cell microvilli are described as being short and brush-like with the microvilli being of the same length. They do not extend far into the taste pore. While one might question the reliability of cell identifications in these studies, both studies relied on immunocytochemistry for gustducin which is a widely-used marker of Type II taste cells. In our preparations of mouse circumvallate taste buds, all cells with short, branched microvilli are Type I cells.

In rat, the microvilli of Type I cells are the longest and extend well into, and often above the taste pore. In the mouse, the Type I microvilli are the shortest and fall into two subtypes: short and tall -- with even the tall microvilli of Type I cells not extending as far outward through the taste pore as either the Type II or Type III cell microvilli. The short-mv Type I cell microvilli that we describe are grossly similar to the microvilli described for the Type II cell of the rat circumvallate taste buds, i.e. short and brush-like with the microvilli being of the same length. They do not extend far into the taste pore. This contrasts with the apical microvilli of mouse circumvallate Type II cells, which consist of a single, stout microvillus that extends through the taste pore into the oral cavity. Rat circumvallate Type III cells possess a single, short, thick, blunt microvillus that does not extend far into the taste pore (Yang, Tabata et al. 2000), while mouse circumvallate Type III cells have a single, long microvillus that projects through the taste pore into the oral cavity. These apparent differences in apical structure may be species differences or might be attributable to different technical approaches or criteria for cell identification.

### Type III Cell mixed synapses

The observation that a significant proportion of Type III cells also possess a mitochondrion with irregular cristae in association with vesicle clusters facing a nerve fiber might be explained in several ways. These Type III cells with nerve-associated mitochondria may represent a functional subset of Type III cells with particular roles in transduction of sour or highly salty tastes. Alternatively, these cells may represent a developmental stage in the natural progression of Type III cells from immature to mature. In either event, the presence of such distinctive mitochondria near the synaptic site may offer a source of ATP for ultimate synaptic release along with other neurotransmitters such as serotonin. Our previous work has shown that purinergic signaling is essential for transmission of Type III cell information (Finger, Danilova et al. 2005, Vandenbeuch, Anderson et al. 2013, Larson, Vandenbeuch et al. 2015) although these cells show no evidence of channel-mediated ATP release as do Type II cells (Taruno, Vingtdeux et al. 2013, Ma, Taruno et al. 2018, Romanov, Lasher et al. 2018). It is possible that Type III cells release ATP from synaptic vesicles, but vesicular loading of ATP requires the vesicular nucleotide transporter (VNUT) which to date, has been localized only to Type II cells (Iwatsuki, Ichikawa et al. 2009).

### Type IV cells

(aka “basal cells”) in taste buds are a population of post-mitotic, relatively immature cells that express sonic hedgehog prior to their ultimate differentiation into the different types of elongate fully mature taste cells (Miura, Kato et al. 2004, Miura, Scott et al. 2014). These cells are generally described as being spherical, based on the nuclear structure, but our analysis of sbfSEM images reveals that the nearly all of the basal cells have a basal process still in contact with the basal lamina perhaps reflecting their relatively recent migration from the more basal position where the proliferative taste bud progenitor cells reside (Miura, Scott et al. 2014). The residual contact with the basal lamina may be important in the final determination of cell fate for this population (Yang, McKee et al. 2011). Similarly, the basal process of Type IV cells intertwines with the basal nerve plexus which likely provides molecular signals crucial for differentiation (Castillo-Azofeifa, Losacco et al. 2017).

The use of sbfSEM 3-D reconstructions greatly enhanced our ability to extract and interpret information from a large data set embracing the full height of a taste bud from basal lamina to taste pore. We believe that this technical approach shows great promise for our future studies in which we plan to elucidate the patterns of synaptic connectivity in taste buds.

## Acknowledgements

The authors would like to thank Logan E. Savidge, Felicia Rodriquez, Daniel Evans who assisted with segmentation of the taste buds and Sue Kinnamon who provided thoughtful discussion and comment on drafts of this manuscript. We also thank the NIDCD of NIH for support of this work.

